# *Phox2b* mutation mediated by *Atoh1* expression impaired respiratory rhythm and ventilatory responses to hypoxia and hypercapnia

**DOI:** 10.1101/2021.08.09.455641

**Authors:** Caroline B. Ferreira, Talita M. Silva, Phelipe E. Silva, Catherine Czeisler, Jose J. Otero, Ana C. Takakura, Thiago S. Moreira

## Abstract

Retrotrapezoid nucleus (RTN) neurons are involved in central chemoreception and respiratory control. Lineage tracing studies demonstrate RTN neurons to be derived from *Phox2b* and *Atoh1* expressing progenitor cells in rhombere 4. *Phox2b* exon 3 mutations cause congenital central hypoventilation syndrome (CCHS), producing an impaired respiratory response to hypercapnia and hypoxia. Our goal was to investigate the extent to which a conditional mutation of *Phox2b* within *Atoh1*-derived cells might affect a) respiratory rhythm; b) ventilatory responses to hypercapnia and hypoxia and c) number of RTN-chemosensitive neurons. Here, we used a transgenic mouse line carrying a conditional *Phox2b^Δ8^* mutation activated by cre-recombinase. We crossed them with *Atoh1^Cre^* mice. Ventilation was measured by whole body plethysmograph during neonate and adult life. In room air, experimental and control groups showed similar basal ventilation; however, *Atoh1^Cre^/Phox2b^Δ8^* increased breath irregularity. The hypercapnia and hypoxia ventilatory responses were impaired in neonates. In contrast, adult mice recovered ventilatory response to hypercapnia, but not to hypoxia. Anatomically, we observed a reduction of the Phox2b^+^/TH^−^ expressing neurons within the RTN region. Our data indicates that conditionally expression of *Phox2b* mutation by *Atoh1* affect development of the RTN neurons and are essential for the activation of breathing under hypoxic and hypercapnia condition, providing new evidence for mechanisms related to CCHS neuropathology.

## Introduction

Breathing is an essential physiological function soon after birth because it can rapidly regulate O_2_ and CO_2_ levels in the blood for the rest of our life. Oxygen is mainly sensed by peripheral chemoreceptors, while CO_2_ can be regulated by central chemoreceptors, and to a lesser extent by peripheral chemoreceptors (Smith et al., 2006; Nattie, 2011; Guyenet, 2014; Guyenet and Bayliss, 2015; Guyenet et al., 2019).

Evidence proposed that an important cluster of excitatory neurons located ventral and immediately caudal to the facial motor nucleus in rodents contains the central chemoreceptors cells. This region was coined in 1989 as the retrotrapezoid nucleus (RTN) (Connely et al., 1990). RTN neurons have a common developmental lineage *(Egr-2, Phox2b, Atoh-1)* (Gaultier and Gallego 2005; Gray, 2008; Champagnat et al., 2011; Heidjen and Zoghbi, 2020) and similar late embryonic (Thoby-Brisson et al. 2009), postnatal (Onimaru et al. 2014) and adult biochemical characteristics (Guyenet and Bayliss, 2015; Guyenet et al., 2009; Guyenet et al., 2019). These neurons are vigorously activated by increases in CO_2_ (Mulkey et al., 2004; Takakura et al., 2006; Moreira et al., 2007; Abbott et al., 2011; Onimaru et al., 2012; Wang et al., 2013; Kumar et al., 2015) and receive inputs from peripheral chemoreceptors (Takakura et al., 2006; 2011; 2014).

*Phox2b* mutations are known to induce disorders in the autonomic control of respiratory system and cause congenital central hypoventilation syndrome (CCHS) (Weese-Mayer et al., 1993; Amiel et al., 2003). CCHS-related *Phox2b* mutations occur in two major categories: a trinucleotide, polyalanine repeat expansion mutations (PARM) or a non- polyalanine repeat expansion mutations (NPARM), that includes missense, nonsense, and frameshift mutations (Patwari et al., 2010; Ramanantsoa and Gallego, 2013; Moreira et al., 2016). The clinical phenotypes of NPARM patients are typically more severe than PARM patients. *Phox2b* polyalanine expansion mutation expression in the RTN neurons results in a blunted response to hypercapnia, but a higher hypoxia ventilatory response (Ramanantsoa et al., 2011). However, the extent to which NPARM expression within RTN neurons could affect the development of breathing and ventilatory responses to hypercapnia and hypoxia remains an open question.

The lack of Phox2b expression of RTN progenitor lineages leads to breathing abnormalities (Ben-Arie et al. 1997; Rose et al., 2009). The *atonal homolog 1* (*Atoh1*, also known as *Math1*) is transiently expressed in RTN neurons (E12.0-P0) and is required for correct RTN progenitor cell migration and differentiation (Dubreuil et al., 2009; Rose et al., 2009; Huang et al., 2012). Studies with *Atoh1*-null mice showed breathing problems shortly after birth and a mislocalized RTN associated with impaired connectivity with the preBötC, suggesting a central mechanism for the respiratory failure and lethality (Ben-Arie et al. 1997; Huang et al., 2012).

In addition, genetic removal of *Atoh1* from *Phox2b* neurons does not impair the CO_2_/H^+^ sensitivity of embryos at E16.5 *in vitro* (Huang et al., 2012) while it does in neonates (Ruffault et al., 2015). Interestingly, conditional mutants that survived showed abnormal hypersensitivity to hypoxia during adult life (Huang et al., 2012). These data suggest a potential developmental interaction between the requirements for *Atoh1* and *Phox2b* in RTN neuron development. To test this hypothesis, we induced the expression of a *Phox2b* non-polyalanine repeat expansion mutation (NPARM) in *Atoh1* progenitor lineage. *Phox2b* NPARM deletions within exon 3 are correlated with severe CCHS phenotype with complete apnoea, profound hypoventilation during sleep and/or cause of post-neonatal infant mortality (Amiel et al., 2003; Weese-Mayer et al., 2010).

We found that mice with mutant *Phox2b* expression conditionally expressed in *Atoh1^Cre^*-derived cells showed a suppressed breath activity to hypoxia and hypercapnia in neonates. We also showed that adult mutant mice presented irregular breathing pattern and the ventilatory response to hypoxia is still compromised while ventilatory response to hypercapnia recovered completely. Despite this, anatomical data showed reduced number of activated neurons by hypercapnia and Phox2b immunoreactivity in the RTN. Together our findings imply that conditionally expression of *Phox2b* mutation by *Atoh1* affect development of the RTN neurons and are essential for the activation of breathing under hypoxic and hypercapnia condition, providing new evidence for mechanisms related to CCHS neuropathology.

## Methods

### 1) Animals

This study was conducted in accordance with the University of Sao Paulo Institutional Animal Care and Use Committee guidelines (protocol number: 3618221019). Our goal was to induce the NPARM mutation in cells giving rise to the CO_2_-sensitive neurons of the RTN. We used a transgenic mouse line with a cre-loxP inducible humanized PHOX2B mutation defined as *Phox2b^Δ8^* and crossed them with Atoh1-cre mice (Nobuta et al., 2015; Alzate-Correa et al., 2020). These animals were bred with Atoh1-cre mice to allow conditional expression of PHOX2B mutant gene (Fig. 1). Genotyping was verified by PCR (REDTaq® ReadyMix™ # R4775, Sigma Aldrich). The primers, genotyping details and strain number of mice used are delineated in Supplementary Table 1.

**Figure 1.**
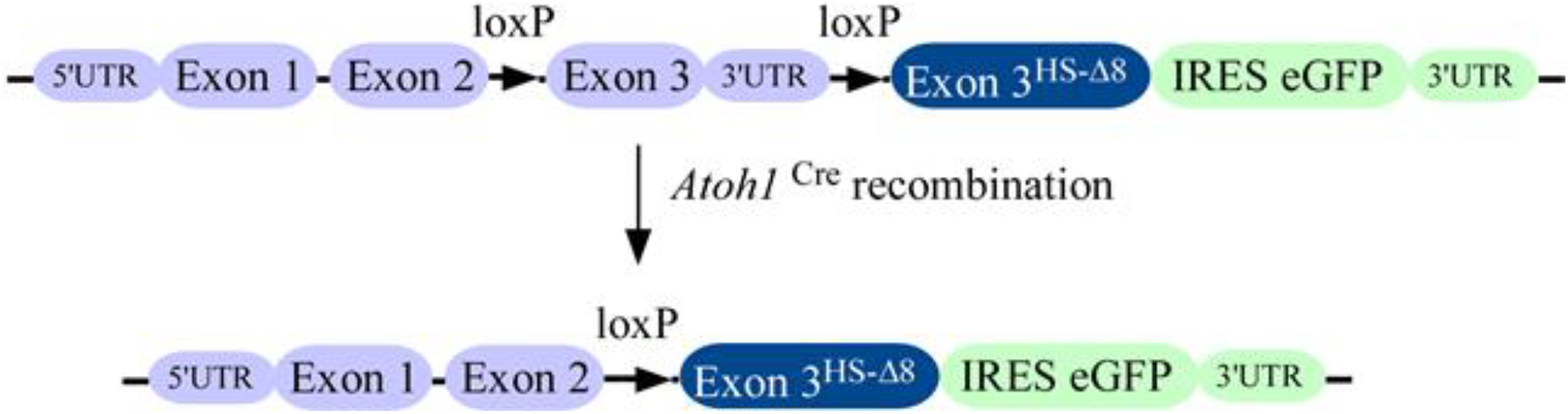
Targeting construct of mouse model. The humanized NPARM PHOX2B mouse mutation is located in exon 3 flanked by loxP sites to allow conditional expression of mutant gene by Atoh1^Cre^ recombinase. Cre-mediated recombination replaces the endogenous murine exon 3 with a human exon 3 harboring an 8-nucleotide deletion (Hs-Δ8).

### 2) Breathing measurements

Breathing variables of neonatal (P1-3) and adult (P30-45) mice from both sexes, were measured noninvasively in unanesthetized and unrestrained using whole-body flow barometric plethysmography (Drorbaugh & Fenn, 1955; Bartlett & Tenney, 1970; Durand et al., 2004). The plethysmograph chamber of neonate had 40 ml and was saturated with water vapor and thermoregulated at 32.5°C and 33.5°C (Durand et al., 2004). The flow rate was set at 0.04 L/min to avoid CO_2_ and water accumulation. Breathing recording in adult mice was done in a larger chamber (500 ml) and flow rate was set as 0.5 L/min. Experiments occurred at 24-26°C room temperature. The animal chamber was connected to a differential pressure transducer and to a preamplifier (FE 141 Spirometer, ADInstruments) to detected pressure oscillations as a result of changes in temperature promoted by ventilation when chamber was completely closed. The signal was digitalized using Power lab system (ADInstrument). The sample rate was set as 1000 Hz and signal were filtered in 0.5 - 20 Hz bandwidth. Volume calibration was performed for each experiment by injecting 2 μL and 100 μL of air into the neonatal and adult chamber, respectively. Breathing variables as breath duration (s), inspiratory time (s), expiratory time (s), tidal volume (V_T_; μl/g), respiratory frequency (f_R_; breaths/min), and ventilation (V_E_; μl/min/g) were analyzed offline using LabChart software (ADInstruments). Tidal volume was calculated as previously described (Patrone et al., 2018). Minute ventilation was defined by the product of breathing frequency and tidal volume. Breath variability was analyzed by inter breath interval (IBI) irregularity and it was defined as IBI irregularity = abs (Ttot (n + 1) – Ttot (n))/ Ttot (n) (Heidjen and Zoghbi, 2018). We also used a nonlinear method of analyses known as Poincare map. This method plots breath duration (Ttot) *vs*. duration of the subsequent breath (Ttot n + 1). We used a total of 100 breaths at rest condition. Next, we calculated the SD1 and SD2 that describes the distribution of the points in the ellipse (Brennan et al. 2001).

To quantify breathing parameters, we first calculated the average of 30 s during a stable condition for each animal during normoxia, hypoxia and hypercapnia. To quantify changes during hypoxia and hypercapnia, we normalized the data to baseline for each animal and then calculate the relative changes expressed as percentage.

### 3) Histology

The mice were deeply anesthetized with isoflurane (5% in 100% O_2_) and heparin were injected intracardially (500 units,) and perfused through the ascending aorta with 20 mL of phosphate-buffered saline (PBS 0.1 M) and with 50 mL of 4% paraformaldehyde (in PBS 0.1 M). The brains were kept overnight immersion in 4% paraformaldehyde and then in a 20% sucrose solution. Brain tissues were sectioned in a coronal plane at 30 μm with a sliding microtome and stored in cryoprotectant solution (20% glycerol plus 30% ethylene glycol in 50 mM phosphate buffer, pH 7.4) at −20°C until histological processing. All histochemical procedures were completed using free-floating sections.

For immunofluorescence, the following primary antibodies were used: a) anti Phox2b (rabbit anti-Phox2b 1:1000; a gift from J.F. Brunet, Ecole Normale Supèrieure, Paris, France); b) anti tyrosine hydroxylase (TH) (mouse anti-TH, 1:1000; Millipore, MA, USA); f) anti fos (rabbit anti-fos, 1:1000; Santa Cruz Biotechnology, CA, USA). All primary antibodies were diluted in PBS containing 2% normal donkey serum (Jackson Immuno Research Laboratories) and 0.3% Triton X-100 and were incubated overnight. Sections were subsequently rinsed in PBS and incubated for 2 hr in an appropriate secondary antibody. The sections were mounted in slides and covered with DPX (Sigma Aldrich, Milwaukee, WI, USA).

### 4) Mapping

A series of three 30 μm transverse sections through the brainstem were examined for each experiment using a Zeiss AxioImager A1 microscope (Carl Zeiss Microimaging, Thornwood, NY). Photographs were taken with a Zeiss MRC camera (resolution 1388 × 1040 pixels). Only cell profiles that included a nucleus were counted and/or mapped bilaterally. Balance and contrast were adjusted to reflect true rendering as much as possible. No other ‘photo retouching’ was performed.

The total number of Phox2b-positive and TH-negative or fos-positive and TH-negative cells in the retrotrapezoide nucleus (RTN: between 5.75 mm and 6.72 mm caudal to bregma level) were plotted as the mean ± S.E.M. (RTN: 12 sections/animal). The neuroanatomical nomenclature employed during experimentation and in this manuscript was defined by the Mouse Brain Atlas from Paxinos and Franklin (2012).

### 5) Experimental protocols

#### Experiment 1: Effect of Phox2b^Δ8^ mutation in Atoh1^Cre^ cells on breathing and chemoreflex activation during neonatal phase

Pups were placed in the plethysmography chambers and acclimated 5 min prior to the experiment. To record breath parameters, the flow was interrupted, and the chamber was closed for 1 min. We recorded a total of 3-5 minutes of ventilation in room air to determine the baseline. To induce chemoreflex challenge, pups were ventilated during 5 minutes in hypercapnia (7% CO_2_, 21% O_2_, balance N_2_) or hypoxia (8% O_2_, balance N_2_) separated by a 10 min of recovery period (room air).

#### Experiment 2: Effect of Phox2b^Δ8^ mutation in Atoh1^Cre^ cells on breathing and chemoreflex activation during adult phase

Adult mice were familiarized during 30 min in 3 consecutive days in the plethysmography chambers. At the day of the breathing recording, animals were acclimated 30 min prior to the experiment. After this acclimation, we recorded 10 minutes in room air breathing to determine the baseline. Animals were then exposed to hypercapnia (7% CO_2_, 21% O_2_, balance N_2_) or hypoxia (8% O_2_, balance N_2_) during 10 min separated by a 20 min of recovery period in room air.

#### Experiment 3: Effect of hypercapnia on fos expression in the RTN neurons after Phox2b^Δ8^ mutation

To investigate whether *Phox2b^Δ8^* mutation change the activation of RTN neurons by hypercapnia, we analyze fos expression in adult mice. Animals were habituated in the plethysmography chambers and ventilated in room air (0.5 L/min) during 3 consecutive days. At the day of experiment, mice were acclimated 1 hour prior to the hypercapnic challenge. Then, animals were exposed to hypercapnia (7% CO_2_, 21% O_2_, balance N_2_) for 45 min. After exposure, mice were ventilated for additional 45 min in room air. Finally, animals were anesthetized and perfused transcardially as described above in “Histology” section. All experiments, were conducted between 9:00 A.M. and 3:00 P.M.

#### Experiment 4: Anatomical changes induced by Phox2b^Δ8^ mutation in the RTN region

To investigate whether *Phox2b^Δ8^* mutation compromised Phox2b expression in the RTN neurons, adult mice were anesthetized and perfused transcardially. Next, tissues were processed by immunohistochemistry to identified Phox2b expression and absence of tyrosine hydroxylase (see details above).

### 6) Statistics

Results are presented as mean + S.E.M. All statistics were performed using GraphPad Prism (version 6, GraphPad Software), with parametric tests used for normally distributed data sets. Details of specific tests are provided in the legend of each figure. The significance level was set as p<0.05.

## Results

### 1) Functional respiratory changes observed in NPARM *Phox2b* mutation in *Atoh1^Cre^* expressing cells

In the following set of experiment, we investigated whether a conditional mutation of *Phox2b*, driven during *Atoh1* expression, in the RTN neurons might affect ventilation during neonatal and adult phase. Given that all *Atoh1^Cre^, Phox2b^Δ8^* mice survived, respiratory parameters were examined between 1-3 and 30-45 post-natal days using whole-body plethysmography. Body weight during neonatal phase was not different between mutation *versus* control littermates (2.08 ± 0.2 g *vs*. 2.27 ± 0.17 g; p = 0.472). In contrast, there were statistic differences in the body weight in animals carrying the mutation during adult life (14.75 ± 0.7 g *vs*. control: 17.3 ± 0.8 g; p = 0.047).

The *Phox2b^Δ8^* mutation mediated by *Atoh1* expressing cells did not affect mean respiratory frequency (neonate mutant: 179 ± 18 bpm *vs.* control: 164 ± 10 bpm, p = 0.679; adult mutant: 231 ± 7 bpm *vs*. control: 218 ± 6 bpm, p = 0.098; Fig. 2A). However, tidal volume was slightly higher during neonatal phase in the mutant mice (12 ± 0.8 μl/g vs. control: 10 ± 0.9 μl/g, p = 0.008; Fig. 2B). On the other hand, despite adult mutant mice showed higher values of tidal volume, it was not statistically different from control littermates (15 ± 1.5 vs. control: 12 ± 0.9 μl/g, p = 0.104; Fig. 2B). Minute ventilation did not change significantly in neonate (mutant: 2.3 ± 0.3 *vs.* control: 1.6 ± 0.2 ml/min/g, p = 0.138; Fig. 2C) and in adult mice (mutant: 3.5 ± 0.3 *vs.* control: 2.7 ± 0.2 ml/min/g, p = 0.104; Fig. 2C). Furthermore, neonates with *Phox2b^Δ8^* mutation did not change inspiratory time (0.114 ± 0.01 *vs.* control: 0.120 ± 0.01 s, p = 0.499; Fig. 2D), expiratory time (0.284 ± 0.04 *vs.* control: 0.318 ± 0.02 s, p = 0.445; Fig. 2E) and total cycle duration (0.40 ± 0.04 s *vs.* control: 0.43 ± 0.03 s; p = 0.557; Fig. 2F) compared to their control littermates. In contrast, during adult phase mice carrying *Phox2b^Δ8^* mutation exhibited a reduction in the inspiratory time (0.081 ± 0.003 s *vs*. control: 0.095 ± 0.002 s; p = 0.026; Fig. 2D) and an increase in the expiratory time (0.21 ± 0.006 s *vs.* control: 0.19 ± 0.005 s; p = 0.025; Fig. 2E), with no change in the total cycle duration (0.293 ± 0.007 s *vs.* control: 0.287 ± 0.006 s; p = 0.629; Figs. 2F).

**Figure 2.**
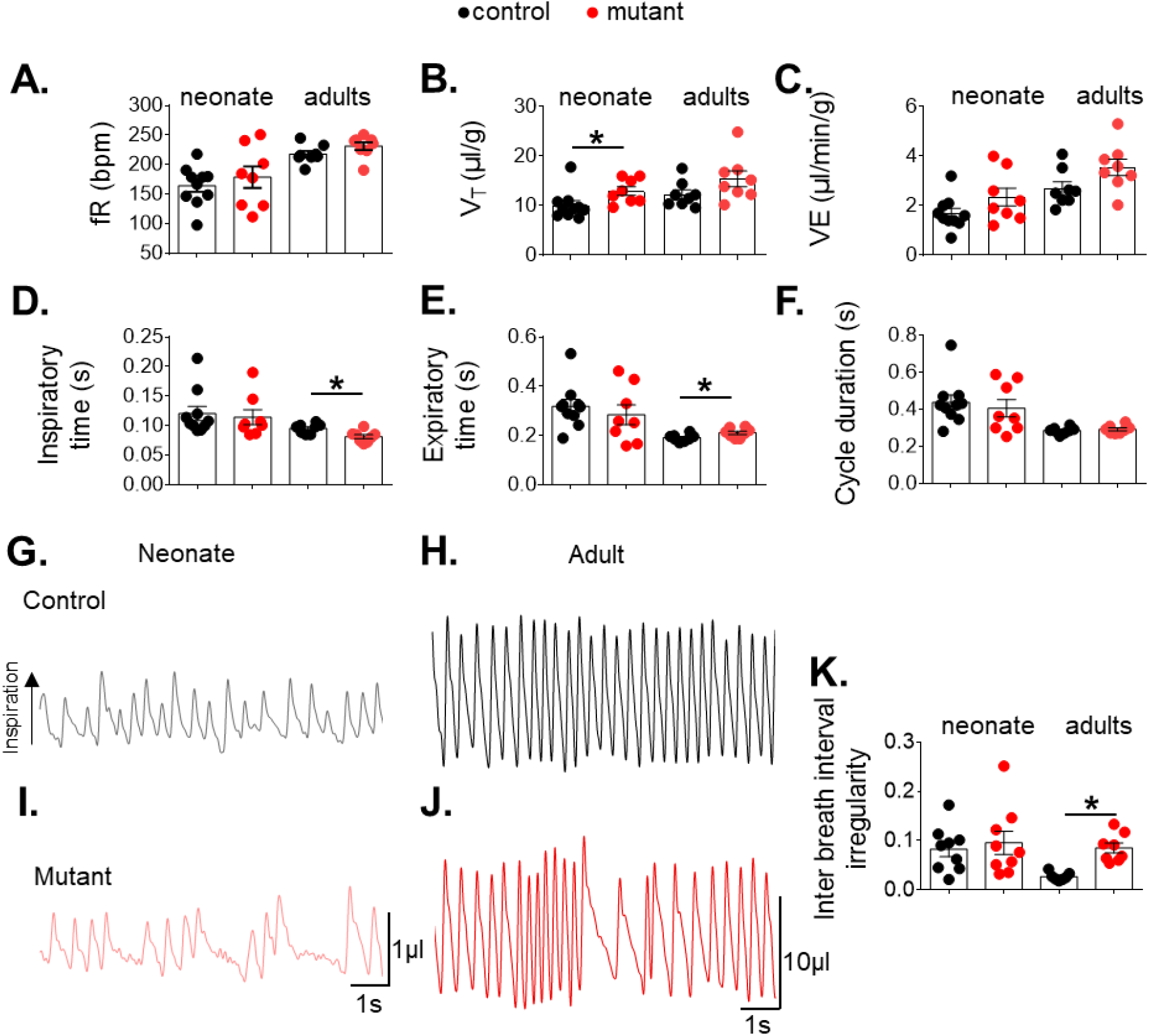
Functional respiratory changes observed in the *Phox2b*^Δ*8*^ mutation in *Atoh1^cre^* cells. Means ± SEM of respiratory frequency (fR; A), tidal volume (V_T_; B), minute ventilation (VE; C), inspiratory time (D), expiratory time (E) and cycle duration (F). (G-J) Plethysmograph recording at rest in control and mutant (*Phox2b*^Δ*8*^;*Atoh1^Cre^*) mice during neonate and adult phase. (K) Means ± SEM of the inter breath interval ((cycle duration (n + 1) - cycle duration(n)/cycle duration(n)). Neonate: control n = 10 and mutant n = 8; Adults: control n = 8 and mutant n = 8. *p < 0.05 *vs.* control from Mann and Whitney U test.

### 2) Selective expression of NPARM *Phox2b* in *Atoh1* cells increased breath irregularity during adult life

Since RTN neurons have been also proposed to participate as generator of respiratory rhythm (Huckstepp et al., 2016; 2018), we next analyzed whether *Phox2b^Δ8^* mutation in RTN neurons might affect breath regularity. Breath-to-breath interval was measured during both neonatal and adult phase. Figure 2 G-J illustrates breathing recording at rest during both neonate and adult phase in controls and mutant mice. There was no change in breath irregularity between mutant and control mice during neonatal phase (0.095 ± 0.02 *vs*. control: 0.082 ± 0.01; p = 0.975; Figs. 2G, I and K). Interestingly, adult mutant mice showed an increase in breath irregularity when compared to their controls (0.085 ± 0.01 *vs.* control: 0.026 ± 0.003; p = 0.0003; Figs. 2H, J and K).

In addition to the time domain analysis, we also used a nonlinear method to investigate breath variability (Li and Nattie, 2006; Patrone et al., 2018; Fernandes-Junior et al., 2018). So, we quantified the distribution of the breath duration using the SD1 and SD2 parameters from the Poincare plots (Fig. 3). *Phox2b^Δ8^* mutation driven by Atoh1cre expressing cells did not affect SD1 and SD2 in neonates (Fig. 3C and D). In the other hand, the adult mutant mice increased the distribution of SD1 and SD2 (Fig. 3E and F). Altogether, these results suggests that NPARM *Phox2b* in RTN neurons affect breath irregularity mainly during adult life.

**Figure 3.**
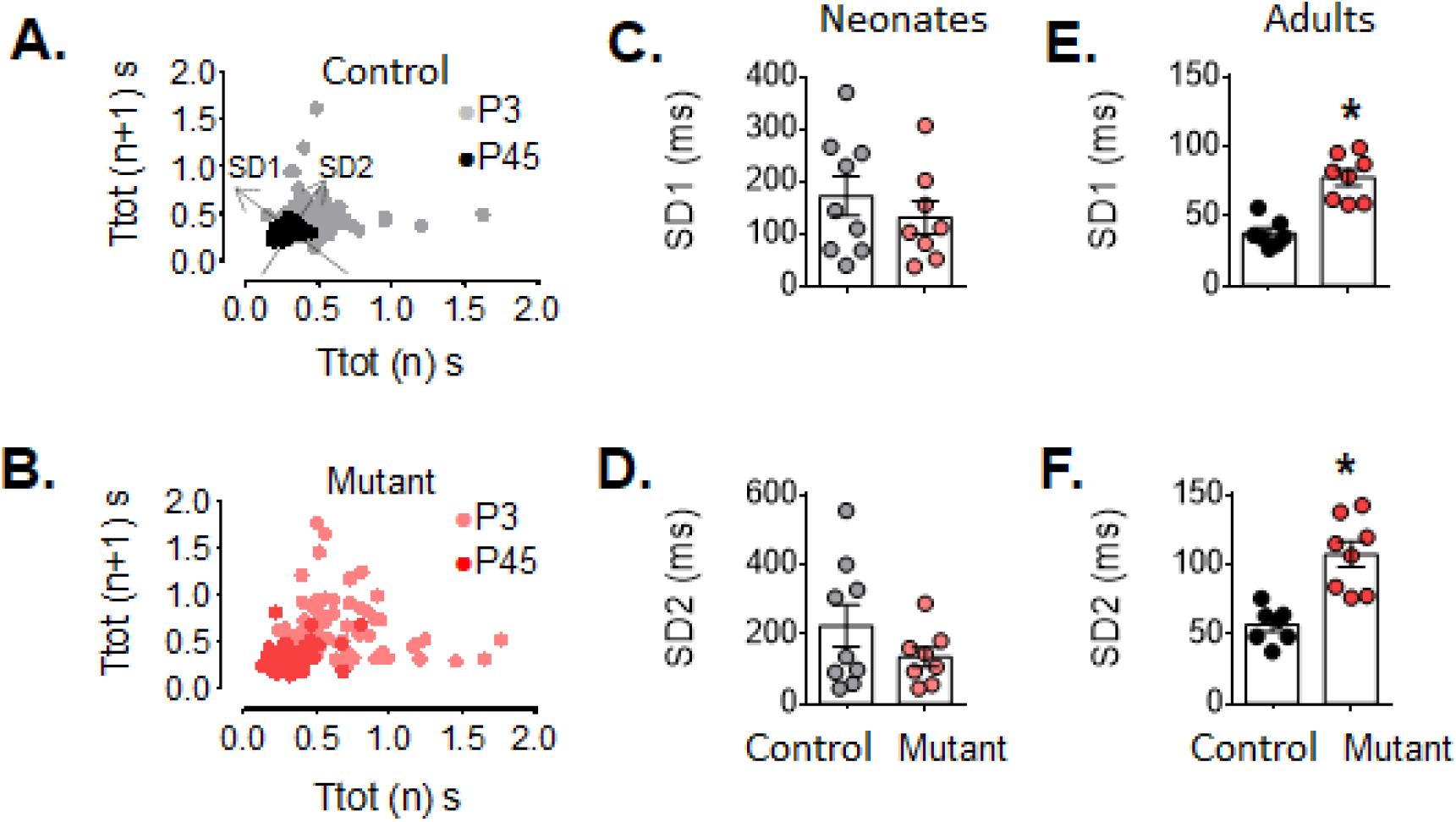
Breath variability increased in mutant adult mice. (A-B) Typical examples of Poincare plot graphs showing SD1 and SD2 from breath duration (Ttot) *vs.* duration of the subsequent breath (Ttot n + 1) in controls and mutant mice (*Phox2b*^Δ*8*^;*Atoh1^Cre^*) during neonatal (P3) and adult (P45) phase. Effect of mutation in SD1 (C) and SD2 (D) during neonatal phase. Effect of mutation in SD1 (E) and SD2 (F) during adult life. Mutation vs. control. *p=0.0003 from Mann and Whitney U test.

### 3) NPARM *Phox2b* in *Atoh1* cells compromised hypoxia and hypercapnia ventilatory response during neonatal phase

A common symptom experienced by patients with the CCHS is an impaired ventilatory response to hypoxemia and hypercapnia (Patwari et al., 2010; Moreira et al., 2016). Therefore, we next explored whether a conditional *Phox2b^Δ8^* within *Atoh1*-derived cells might impair ventilatory response to hypoxia during the first days of life. Figures 4A-B illustrates an example of breathing responses to 8% O_2_ in a control and mutant mice at day 3 after birth. We monitored baseline ventilation while neonates were breathing room air followed by 5 min of 8% O_2_. We choose to analyze the first minute of hypoxic exposure because longer than 5 min of low O_2_ exposure lowers body temperature in mice (Kline et al. 1998). Figures 4E-G illustrates normalized changes in the respiratory frequency, tidal volume, and minute ventilation at the first minute of hypoxia. As expected, when exposed to hypoxia control littermates increased respiratory frequency around 40% (from: 100 ± 6% to: 139 ± 5%; p = 0.0016; Fig. 4E) and ≈ 45% in the tidal volume (from: 100 ± 9% to: 146 ± 15%; p = 0.0058; Fig. 4F), resulting in a significant increase in minute ventilation (from: 100 ± 12% to: 205 ± 24%; p=0.0076; Fig. 4G). On the other hand, the increase in respiratory rate during hypoxia in mutant pups were not different from normoxia (from 100 ± 10 to 123 ± 8%; p = 0.26; Fig. 4E). Furthermore, neonate mutant mice failed to increase tidal volume (from 100 ± 6 to 105 ± 9%; p = 0.91; Fig. 4F). As a result, there was no statistic difference in minute ventilation in response to hypoxia (from 100 ± 15 to 128 ± 11%; p = 0.37; Figs. 4G).

**Figure 4.**
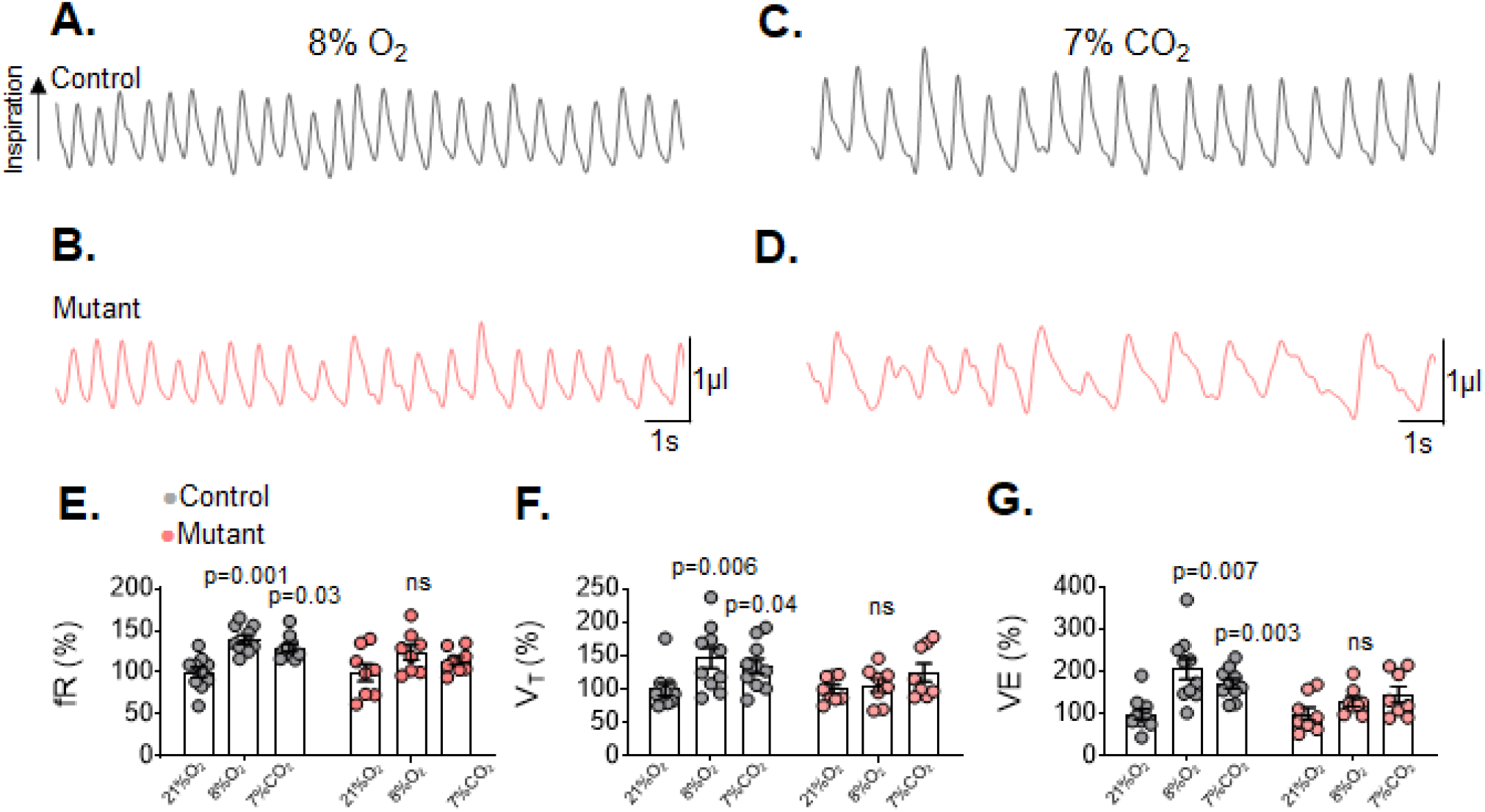
*Phox2b*^Δ*8*^ mutation impaired ventilatory responses to hypoxia and hypercapnia in neonates. Plethysmograph breathing traces showing ventilation in a control and mutant (*Phox2b*^Δ*8*^;*Atoh1*^Cre^) mice while ventilated with 8% O_2_ (A and B) and 7% CO_2_ (C and D). E) Changes in respiratory frequency (f_R_; interaction: F (2, 32) = 0.8, p = 0.455; effect of mutation F (1, 16) = 4.3, p = 0.052; effect of hypoxia and hypercapnia: F (2, 32) = 10, p = 0.0003). F) Changes in tidal volume (V_T_; interaction: F (2, 32) = 1.9, p = 0.162; effect of mutation: F (1, 16) = 2.4, p = 0.138); effect of hypoxia and hypercapnia: F (2, 32) = 4.4, p = 0.019). G) Changes in minute ventilation (V_E_; interaction: F (2, 32) = 2.102, p = 0.138; effect of mutation: F (1, 16) = 4.482, p = 0.0503); effect of hypoxia and hypercapnia: F (2, 32) = 12.89, p < 0.0001); ANOVA 2-way Dunnett’s multiple comparisons test.

Our next goal was to investigate whether a conditional *Phox2b^Δ8^* mutation affects ventilatory response to hypercapnia. Figure 4C-D illustrates a typical respiratory trace from the same cre-negative and neonate mutant mice but now ventilated with 7% of CO_2_. Control pups increased respiratory frequency by approximately 30% during hypercapnia (p = 0.032; Fig. 4E) when compared to normoxia. In addition, tidal volume changed significantly from 100% ± 9% to 134% ± 11% (p = 0.042; Fig. 4E). Therefore, there was an increase of 70% in the minute ventilation (p = 0.0033; Fig. 4G). In contrast, *Phox2b^Δ8^* mutation failed to increase respiratory frequency (from: 100% ± 10% to: 113% ± 5%; p = 0.36; Fig. 4E) and tidal volume (from: 100% ± 6% to: 124% ± 13%; p = 0.22; Fig. 4F) in response to hypercapnia. Therefore, the minute ventilation did not change significantly from normoxia (from: 100% ± 15% to: 144% ± 18%; p = 0.22; Fig. 4G). It suggests that *Phox2b^Δ8^* mutation driven by *Atoh1* cells in RTN neurons affect ventilatory responses to hypoxia and hypercapnia during neonatal phase.

### 4) Ventilatory response to hypoxia is still compromised in the *Atoh1^Cre^, Phox2b^Δ8^* adult phase

The hypoxic ventilatory responses were also explored during adult phase. Figures 5A-B illustrate an example in a control and mutant adult mice. Baseline ventilation was monitored while animals were breathing room air followed by 10 min of hypoxic challenge. Figure 5C-E demonstrates that the ventilatory responses were partially recovered in adult mutant mice. Both mutant and control, showed a biphasic response characteristic of hypoxic stimulus. However, the magnitude of the ventilatory responses were reduced in the mutant adult animals. As demonstrated in Figure 5C, respiratory frequency had a peak in the first minute for both control (from: 100 ± 4 to: 146 ± 5; p=0.0009) and mutant mice (from: 100 ± 5 to: 154 ± 10; p=0.032). After the third minute, the tidal volume reduced mainly in the mutant across the 10 min of stimulus (Fig. 5D). As a result, the mutants had a lower minute ventilation (Fig. 5E). These results demonstrates that mutant mice had an impaired ventilatory response to hypoxia in the adult phase.

**Figure 5.**
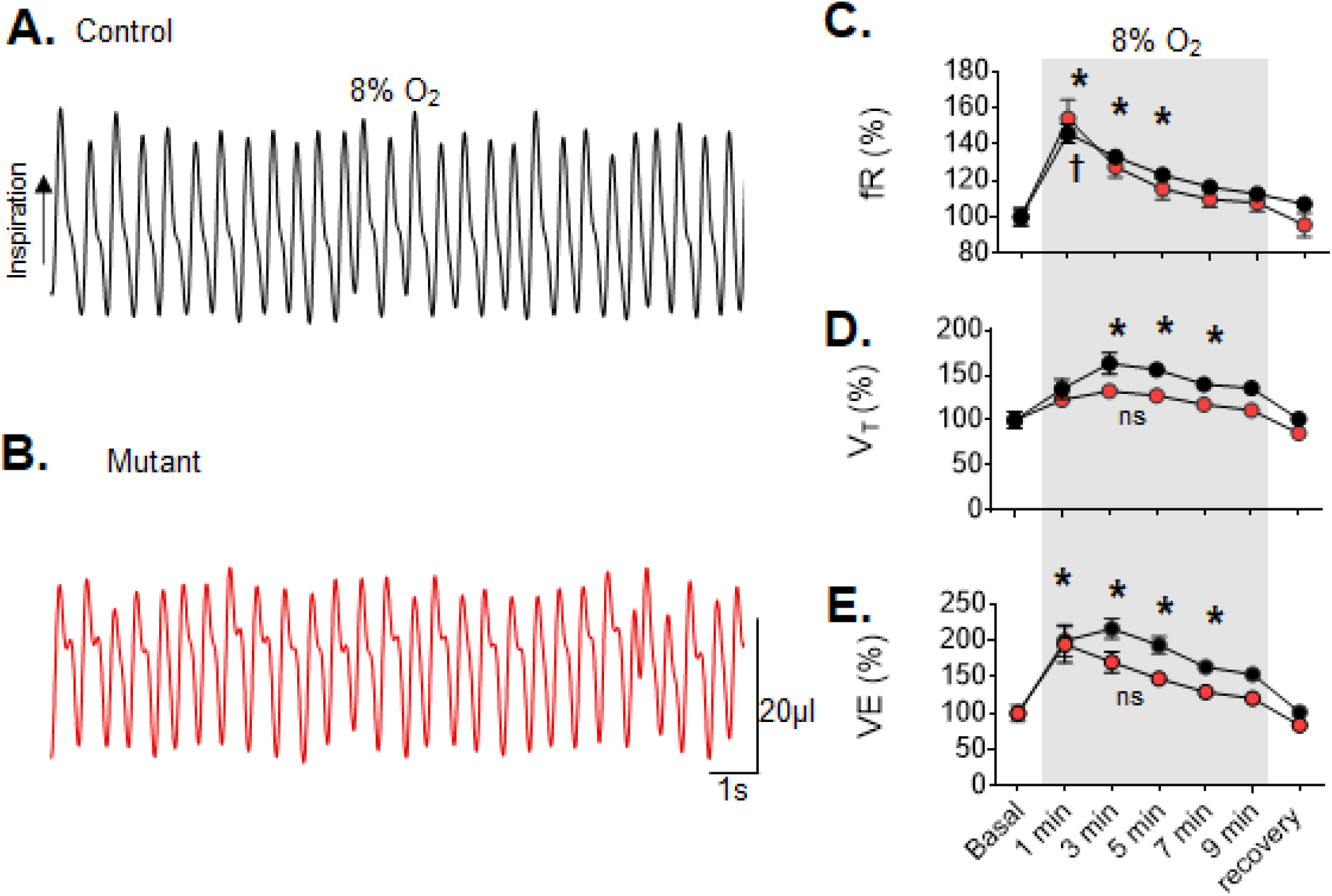
Ventilatory responses induced by hypoxia are partially recovered during adult life in mutant mice. Plethysmograph traces showing ventilation in a control (A) and mutant (*Phox2b*^Δ*8*^;*Atoh1*^Cre^; B) mice during hypoxia (FiO_2_ = 8%). Mean ± SEM from relative changes of respiratory frequency (C; fR; interaction: F (6, 84) = 0.972, p = 0.45; effect of mutation F (1, 14) = 0.311, p = 0.58; effect of hypoxia: F (2.082, 29.15) = 69.77, p < 0.0001)), tidal volume (D; V_T_; interaction: F (6, 84) = 0.972, p = 0.45; effect of mutation F (1, 14) = 0.311, p = 0.58; effect of hypoxia: F (2.082, 29.15) = 69.77, p < 0.0001)) and minute ventilation (E; V_E_; interaction: F (6, 84) = 1.195, p = 0.31; effect of mutation F (1, 14) = 8.218, p = 0.0124; effect of hypoxia: F (2.227, 31.18) = 23.92, p < 0.0001) in response to hypoxia. *p < 0.05 *vs.* control in 21% O_2_. †p < 0.05 *vs.* mutant in 21% O_2_. ANOVA 2-way Dunnett’s multiple comparisons test.

### 5) *Atoh1^cre^, Phox2b^Δ8^* recovered hypercapnia ventilatory response during adult phase

The next step was to investigate whether ventilatory response to hypercapnia were impaired during adult life. At rest, ventilation was recorded in room air followed by 10 min of hypercapnic challenge. A striking effect was that mutation completely recovered ventilatory response in adults carrying *Atoh1^Cre^, Phox2b^Δ8^* mutation (Figs. 6A-E). Figure 6A-B shows an example of breathing while a control and mutant mice were ventilated with 7% of CO_2_. Both groups, markedly increased minute ventilation due an increase in tidal volume and respiratory frequency (Figs. 6C-E). These results demonstrated that adult mutant mice recovered their ventilatory response to hypercapnia.

**Figure 6.**
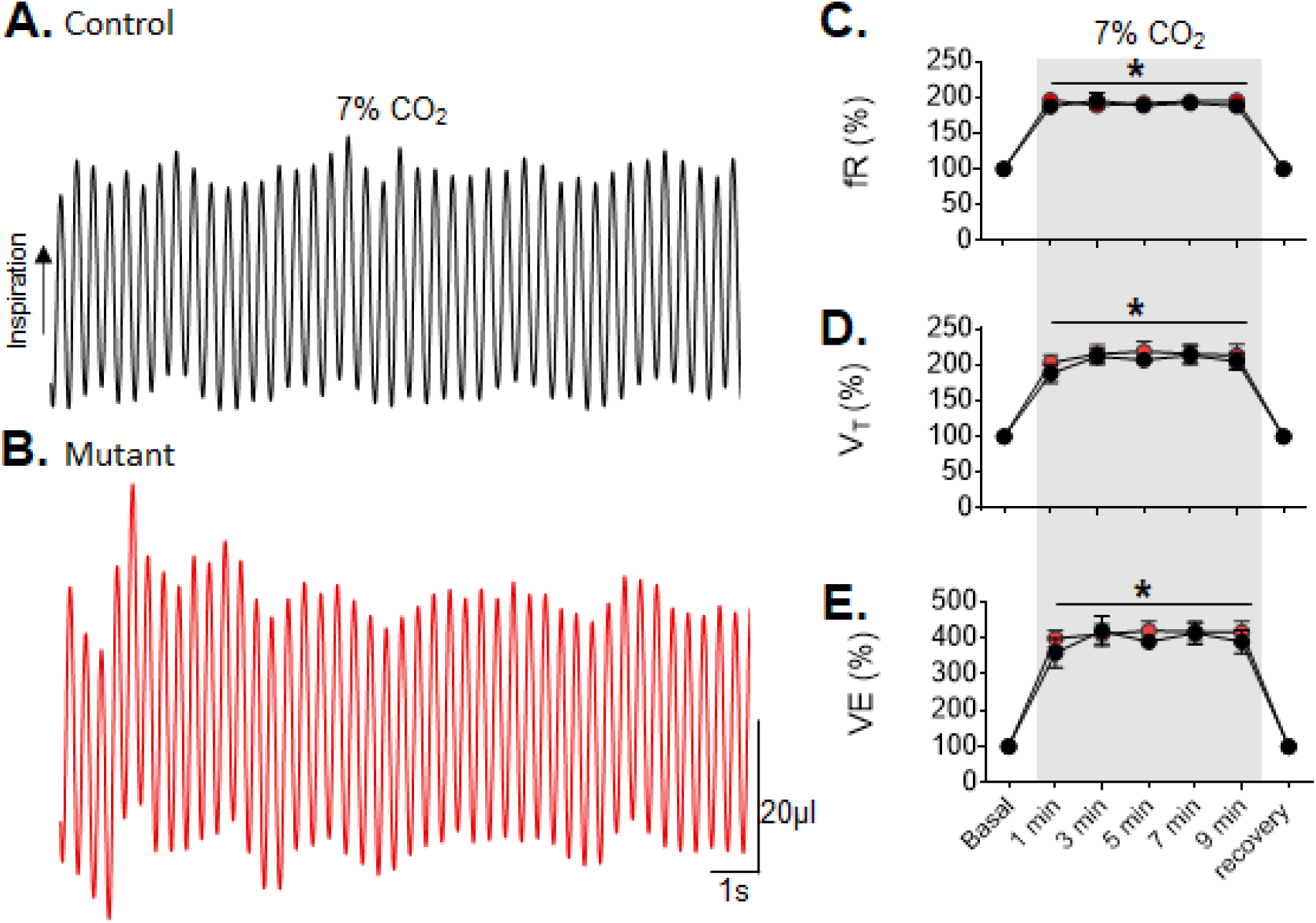
Adult *Phox2b*^Δ*8*^;*Atoh1^Cre^* mice recovered ventilatory responses to hypercapnia. Plethysmograph traces showing ventilation in a control (A) and mutant (*Phox2b*^Δ*8*^;*Atoh1^Cre^*; B) mice while breathing 7% of CO_2_. Mean ± SEM from normalized changes in the respiratory frequency (C; fR; interaction: F (6, 84) = 0.56, p = 0.76; effect of mutation F (1, 14) = 0.41, p = 0.53; effect of hypercapnia: F (6, 84) = 155.48, p < 0.0001)), tidal volume (D; VT; interaction: F (6, 84) = 0.22, p = 0.96; effect of mutation F (1, 14) = 0.31, p = 0.58; effect of hypercapnia: F (2.082, 29.15) = 69.77, p < 0.0001)) and minute ventilation (E; VE; interaction: F (6, 84) = 0.33, p = 0.91; effect of mutation F (1, 14) = 0.37, p = 0.54; effect of hypercapnia: F (2.206, 30.88) = 86.85, p < 0.0001) in response to hypercapnia. *p ≤ 0.0005 *vs.* 21% O_2_ for both control and mutation group. ANOVA 2-way Dunnett’s multiple comparisons test.

### 6) NPARM *Phox2b* in *Atoh1* cells reduced the activation of RTN neurons by hypercapnia

Since mutant mice recovered the ventilatory response to hypercapnia in adulthood, the next set of experiments were done to explore whether the RTN neurons might be activated in the *Atoh1cre, Phox2b^Δ8^* animals using the fos as a reporter of cell activation. Hypercapnia induces *fos* expression in the rodent RTN (Sato et al. 1992; Teppema et al., 1994; Fortuna et al., 2009; Kumar et al., 2015; Shi et al., 2021). Thus, we compared fos expression by the RTN neurons in the control and mutant mice while exposing to hypercapnia. In the *cre* negative mice, RTN neurons lacking TH immunoreactive showed elevated fos immunoreactivity when exposed to hypercapnia. In contrast, the number of fos-activated neurons in the RTN significantly reduced in the mutant mice (37 ± 8 *vs.* control: 86 ± 19 neurons; p = 0.047; Figs. 7A-C). It shows that despite adult mutant mice recovered the ventilatory response to hypercapnia the number of RTN neurons activated by hypercapnia were reduced.

**Figure 7.**
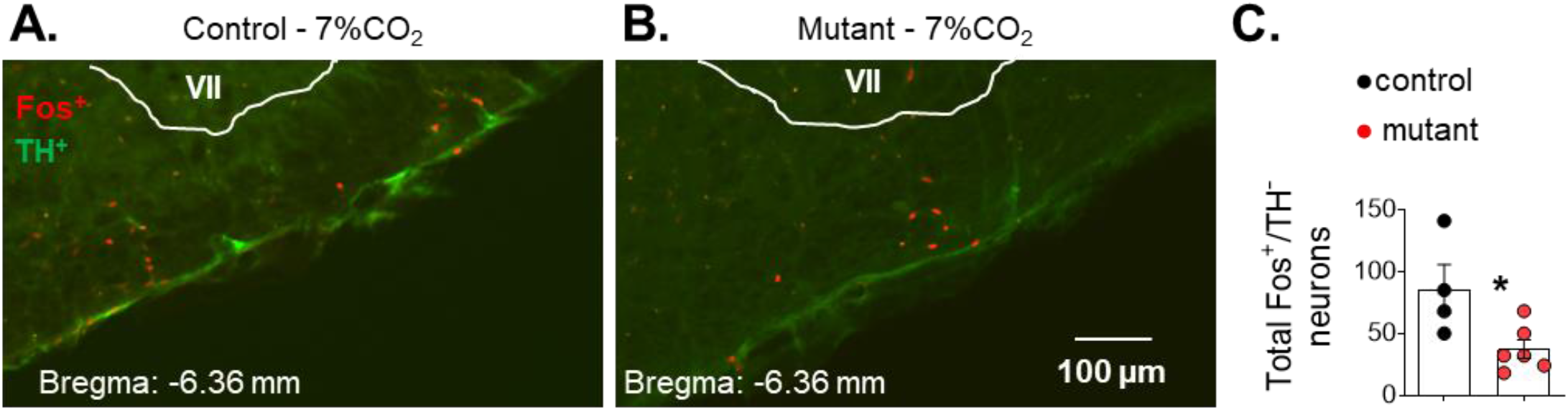
Fos-activated neurons in the RTN in response to hypercapnia are reduced in mutant adult mice. Photomicrographs from a control (A) and mutant (*Phox2b*^Δ*8*^;*Atoh1^Cre^*; B) mice exposed to 7% of CO_2_. (C) Mean ± SEM of neurons expressing fos in the absence of tyrosine hydroxylase (TH). Control n = 4 and mutant n = 6. VII: facial nucleus. *p = 0.047 *vs*. control 7% CO_2_. Mann Whitney test.

### 7) NPARM *Phox2b* in *Atoh1* cells reduced Phox2b immunoreactivity in the RTN neurons

The CO_2_-sensitive cells of the RTN belong to a neuronal group with a well-defined phenotype characterized by the presence of Phox2b-ir and the absence of tyrosine hydroxylase (TH) (Stornetta et al., 2006). The transcription factor *Atoh1* is also expressed in the RTN neurons during developmental stage (van der Heijden and Zoghbi; 2020). This transcription factor has a key role in the hindbrain development and was used as *cre* driver to induce recombination in cells giving rise to the CO_2_ sensitive neurons of the RTN (Wang et al., 2005; Rose et al., 2009). Therefore, to describe and understand *Atoh1cre, Phox2b^Δ8^* neuropathology, we investigated whether the mutation induces anatomical changes in the RTN neurons. For this purpose, we investigated the number of Phox2b-expressing neurons at the level of RTN in the adult mice. Figures 8 demonstrates typical photomicrographs from different levels of RTN in a cre-negative (Figs. 8A-C) and mutant mouse (Figs. 8D-F). Phox2b^+^/TH^−^ expression reduced significantly in the adult *Atoh1cre, Phox2b^Δ8^* mice when compared to respective control (143 ± 41 *vs.* control: 266 ± 16; p = 0.0317; Fig. 8G).

**Figure 8.**
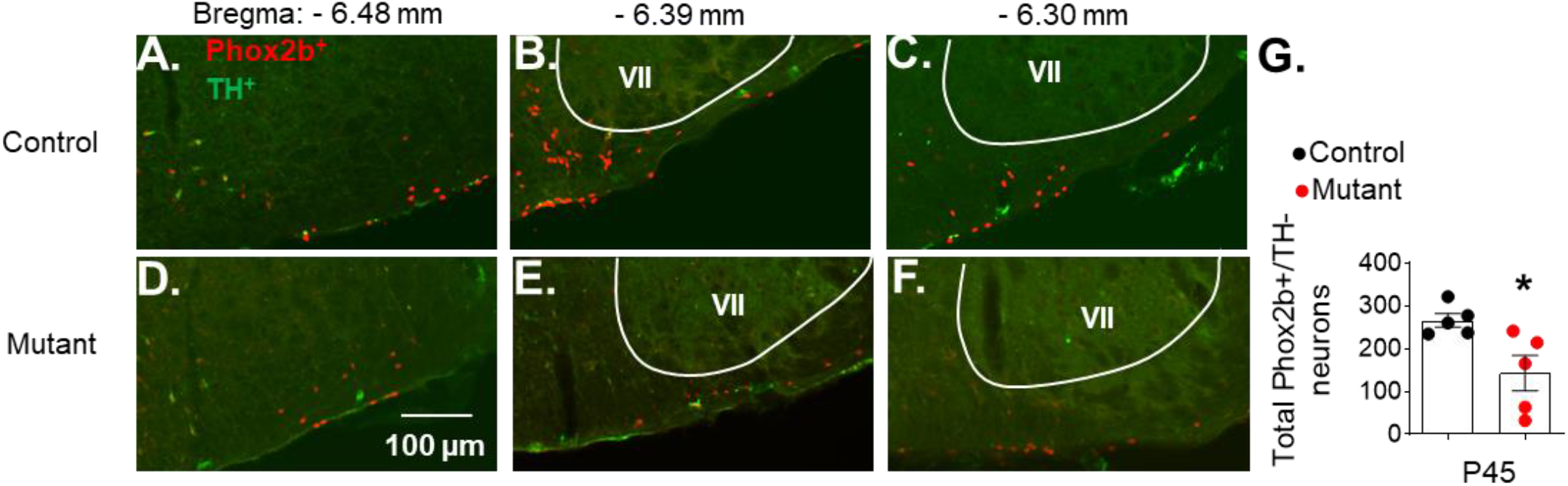
Adult mutant mice reduced Phox2b expression in the RTN neurons. Photomicrographs from the three different levels of RTN in a control (A-C) and mutant (*Phox2b*^Δ*8*^;*Atoh1^Cre^*; D-F) mice. (G) Mean ± SEM of total neurons expressing Phox2b in the absence of tyrosine hydroxylase (TH). Control n = 5 and mutant n = 5. VII: facial nucleus. *p = 0.0317 *vs*. control. Mann Whitney test.

## Discussion

The designation of specific behaviors to small clusters of neurons represents a premier challenge to the neuroscience community. A premier example of this challenge is the RTN chemoreceptor neurons, which despite having fewer than 800 neurons bilaterally shows significant regulation of breathing. The RTN neurons are located in the ventral medullary surface and have been described to be involved in breathing regulation by CO_2_/H_+_ (Li and Nattie, 2002; Mulkey et al., 2004; Takakura et al., 2006; Stronetta et al., 2006; Abbott et al., 2009; Huckstepp et al., 2010; Marina et al., 2010; Nattie, 2011; Guyenet and Bayliss, 2015). In the present study, we reexamined the role of RTN on breathing activity and chemoreception process using a genetic background based on prior evidence that RTN neurons express and require the *Phox2b* and *Atoh1* transcription factors (Dubreuil et al., 2009; Rose et al., 2009; Ramanantsoa et al., 2011). We demonstrated that mutation in the *Phox2b* located in *Atoh1^Cre^*-expressing cells showed i) irregular breathing pattern, ii) reduced hypoxia ventilatory response in neonate and adult, iii) adult mutant mice recovered the hypercapnia ventilatory response despite a reduction of fos-activated neurons in the RTN, and iv) a significant reduction of Phox2b-expressing cells within RTN region in adults. Together our findings imply that mutant *Phox2b* expression in the *Atoh1* neurons of the RTN are obligatory elements of the circuitry for breathing regulation by chemosensory control of breathing in mice.

### Role of *Phox2b*^+^/*Atoh-1*^+^ RTN neurons on respiratory chemoreception

Our physiological data showed a reduction in the hypoxia and hypercapnia ventilatory response in neonate mice. Our data in both adults and neonates are partially in line with other findings in the literature generated from PARM Phox2b mutant mice (Dubreuil et al., 2009; Marina et al., 2010; Ramanantsoa et al., 2011; Takakura et al., 2014; Ruffault et al., 2015). Phox2b^+^ expressing RTN neurons are important CO_2_ sensors in the brain and receive chemosensory inputs from other cells in the respiratory column in the brainstem (Rosin et al., 2006; Guyenet et al., 2005). Briefly, RTN neurons are i) sensitive to small changes in CO_2_/H^+^ (Mulkey et al., 2004; Onimaru et al., 2008; Wang et al., 2013); ii) in close opposition to numerous capillaries (Onimaru et al., 2012; Hawkins et al., 2017; Cleary et al., 2020), classifying these neurons to be critical to sense CO_2_ in the blood; iii) RTN neurons also receive afferents from many brainstem sites that contain putative chemosensors (Rosin et al., 2006); iv) respond with depolarization to activation of nearby acid-sensitive astrocytes (Gourine et al., 2010; Wenker et al., 2010; 2012), and v) receive excitatory connections from the carotid bodies (CBs) (Takakura et al., 2006). Therefore, the main question is: what mechanisms enable the mutants to breathe and to presumably maintain normal blood P_CO2_ in a condition where the chemosensors neurons in the RTN were reduced? Possibilities include strengthening of the CO_2_-responsive input from the CBs or compensation by one or the other of the multiple chemoreceptor sites in the brain (Nattie, 2011).

The first possibility is that carotid body compensates for the CO_2_-drive to breathe and then through nucleus of the solitary tract (NTS) activates the respiratory column to maintain breathing activity. The plausible explanation emerges by considering that RTN neurons are strongly activated by carotid body stimulation and provide powerful excitatory input to the respiratory column (Takakura et al., 2006). They may thus be obligatory intermediates for relaying the CO_2_ response when we have a loss of RTN neurons. The second possibility is that the RTN is not an obligatory site for central chemoreceptors. Candidate chemoreceptor sites are serotonergic neurons that have been reported to be pH-sensitive (Wang and Richerson, 1999; Corcoran et al., 2009), the noradrenergic neurons located in the locus coeruleus (Biancardi et al., 2008) and glial cells (Gourine et al., 2010; Wenker et al., 2010; 2012; Sobrinho et al., 2014).

The results of previous loss-of-function experiments to assess the role played by RTN neurons in the chemoreflex are not entirely conclusive. In previous work, we evaluated the chemoreflex in which subsets of Phox2b-expressing neurons in the RTN were lesioned using toxin or pharmacological tools (Takakura et al., 2006; 2008; 2013; 2014). Bilateral lesions of the neurokinin1 receptor-expressing neurons in the RTN region by injection of saporin conjugated to a substance P or injection tof he GABA-A agonist muscimol have produced a reduction of the hypercapnic ventilatory response (Nattie and Li, 2002; Takakura et al., 2008; 2013; 2014). Unfortunately, these experiments lack specificity, and the extent of the lesion or inhibition is very difficult to control. Using a more selective approach, Marina and colleagues (2010) used a pharmacogenetic tool to silence RTN neurons. Rats that had received injection of lentivirus vector expressing the allatostatin receptor from PRSx8 promoter reduced the hypercapnic ventilatory response after administration of allatostatin. However, the PRSx8 promoter used targets the neurons in the rostral aspect of the ventrolateral medulla, including RTN, C1 adrenergic and A5 noradrenergic neurons (Stornetta et al., 2006; Abbott et al., 2013; Burke et al., 2014; Malheiros-Lima et al., 2018; 2020). Finally, using genetic backgrounds, RTN neurons were deleted and the chemoreflex lost, but the effect could not be entirely attributed only to the RTN (Dubreuil et al., 2008, 2009; Ramanantsoa et al., 2011). A similar response of the hypercapnic ventilatory response described in adult *Phox2b^Cre^*;*Atoh1*^lox/lox^ mutants mice was previously demonstrated (Ruffault et al., 2015). The recovery of the CO_2_ response in adults could be due to late compensation by residual RTN neurons, to activation of peripheral chemoreceptor input that bypasses the RTN (Basting et al., 2015) or to some of the multiple chemosensors sites as described before (Nattie, 2011).

Hypoxic ventilatory response emerges from a physiological reflex of the already established notion that ventrolateral brainstem respiratory neurons are excited by peripheral chemoreceptors via a direct glutamatergic input from commissural NTS (Guyenet, 2014). Besides the di-synaptic excitatory pathway from commissural NTS to RVLM, we also know that we have a relay via the chemosensitive neurons of the RTN (secondary input) (Takakura et al., 2006). Therefore, the compromised ventilatory response to hypoxia in the present study can be explained by the fact that this pathway was affected by the conditional mutation of RTN neurons during *Atoh1* expression.

The open question that needs to be investigated is by which mechanism *Phox2b* mutation mediated by *Atoh1* can change chemosensory control of breathing. Such mechanisms may be involved in selective loss of neurons, disorganized respiratory circuits, and likely contributes to the irregular breathing pattern and severe apneic phenotype. In addition, it is important to investigate whether the recovery of the hypercapnia ventilatory response in adults could be due to a plasticity of remaining RTN neurons and/or to other chemosensitivity areas assuming this function.

## Conclusion

A consensus about the sites and circuits that underlie the chemosensory control of breathing has yet to emerge. Our data established that the *Phox2b*^+^/*Atoh1*^+^ RTN neurons, are involved in the pathway of the chemosensory control of breathing. In both neonate and adult, they are necessary for the hypoxia ventilatory response. Irregular breathing is characteristic for preterm infants and not uncommon in babies that are born at term (Gaultier and Gallego, 2005). The key role of the RTN for breathing regulation at birth that we show here suggests that defective development or immaturity of the human equivalent, apart from causing CCHS (Amiel et al., 2003), could also underlie more common respiratory problems in the newborn period.

## Acknowledgement

This research work was supported by public funding from São Paulo Research Foundation (FAPESP) (Grants: 2019/01236-4 to ACT and 2015/23376-1 to TSM); and by funds from FAPESP fellowship (2017/12678-2 to TMS and 2019/20990-1 to PES), Conselho Nacional de Desenvolvimento Científico e Tecnológico (CNPq) grant (408647/2018-3 to ACT) and fellowships (302334/2019-0 to TSM and 302288/2019-8 to ACT) and NHLBI/NIH (Grant: R01HL132355 to CMC and JJO). This study was financed in part by the Coordenação de Aperfeiçoamento de Pessoal de Nível Superior-Brasil (CAPES) - Finance Code 001.

---

**Supplementary Table 1:**
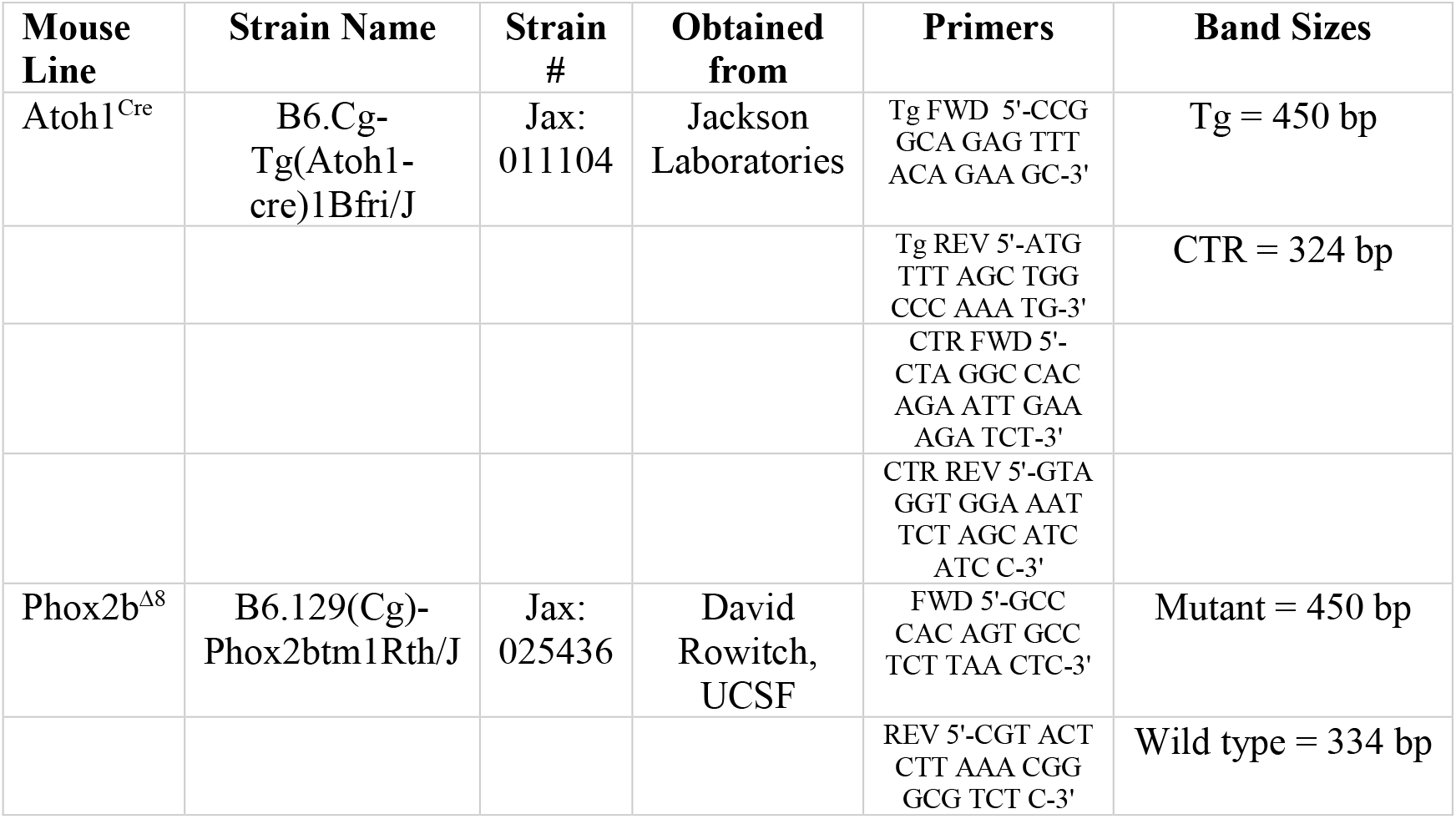
Genotyping primers.

